# Serine proteinase inhibitors from *Nicotiana benthamiana*, a non-preferred host plant, inhibit the growth of *Myzus persicae* (green peach aphid)

**DOI:** 10.1101/2023.05.16.540980

**Authors:** Honglin Feng, Georg Jander

## Abstract

The green peach aphid (*Myzus persicae*) is a severe agricultural crop pest that has developed resistance to most current control methods, requiring the urgent development of novel strategies. Plant proteinase inhibitors (PINs) are small proteins that protect plants against pathogens and/or herbivores, likely by preventing efficient protein digestion. We identified 67 protease genes in the transcriptomes of three *M. persicae* lineages (USDA-Red, G002, and G006). Comparison of gene expression levels in aphid guts and whole aphids showed that several proteases, including a highly expressed serine protease, are significantly overexpressed in the guts. Furthermore, we identified three genes encoding serine protease inhibitors (*SerPIN-II1*, *2*, and *3*) in *Nicotiana benthamiana*, which is a non-preferred host for *M. persicae*. Using virus-induced gene silencing (VIGS) with a tobacco rattle virus (TRV) vector and overexpression with a turnip mosaic virus (TuMV) vector, we demonstrated that *N. benthamiana SerPIN-II1* and *SerPIN-II2* cause reduced survival and growth, but do not affect aphid protein content. Similarly, *SerPIN-II3* overexpression reduced survival and growth, and *serpin-II3* knockout mutations, which we generated using CRISPR/Cas9, increased survival and growth. Whereas protein content was significantly increased in aphids fed on *SerPIN-II3* overexpressing plants, it was decreased in aphids fed on *serpin-II3* mutants. Together, our results show that three PIN-IIs from *N. benthamiana*, a non-preferred host plant, effectively inhibit *M. persicae* survival and growth, thereby representing a new resource for the development of aphid-resistant crop plants.

## Introduction

Aphids are severe agricultural pests that cause damage to a wide range of crops, not only by direct feeding, but also more seriously by spreading plant viruses, promoting growth of sooty mold, and blocking photosynthesis (Jayasinghe et al., 2021). The green peach aphid, *Myzus persicae*, is a broad generalist that can colonize hundreds of host plant species from 40 families, including woody hosts (*e.g.*, peach), vegetables (*e.g.*, cabbage, cucumber, eggplant, potato, and tomato), ornamental flower crops, and field crops (*e.g.*, sunflower and tobacco) (Blackman & Eastop, 2008). *Myzus persicae* is considered a serious vector of plant viruses and can transmit hundreds of viruses across different host plants, including viruses from the Luteoviridae, Potyviridae, and Nanoviridae families, through both non-persistent and persistent transmission (Gadhave et al., 2020; Hogenhout et al., 2008). Another major concern with *M. persicae* is that the rapid development of insecticide resistance. This species has developed resistance to more than seventy insecticides (Vasquez, 1995), and at least seven genetically distinct mechanisms of resistance have been characterized (Bass et al., 2014). Therefore, novel strategies are needed for controlling *M. persicae* infestations.

*Nicotiana benthamiana*, a wild tobacco species, is commonly used as a model for plant molecular biology research. Virus-induced gene silencing (VIGS) with tobacco rattle virus (TRV) can reduce gene expression in both the host plants and herbivorous insects (Bachan & Dinesh-Kumar, 2012; Feng et al., 2023; Feng & Jander, 2022), and it is possible to implement tissue culture-independent germline editing in *N. benthamiana* using CRISPR/Cas9 (Čermák et al., 2017; Cody et al., 2022; Ellison et al., 2020). As a non-preferred host plant for many tobacco-feeding generalist herbivores, *N. benthamiana* has been used to investigate different mechanisms of plant-insect defense and counter-defense, including aphid effector proteins (*e.g.*, Mp10, Mp42 (Rodriguez et al., 2014), and Mp55 (Elzinga et al., 2014)) and acylsugars as a plant chemical defense (Feng et al., 2021; Wang et al., 2022).

Proteinase inhibitors (PINs) are among the most abundant classes of proteins and are found throughout all life forms, including microorganisms, insects, plants, and animals (Ryan, 1989). PINs have two main functions in plants: 1) They prevent uncontrolled endogenous proteolysis, which is crucial for physiological processes such as programmed cell death (Birk, 2003; Lampl et al., 2013; Solomon et al., 1999); and 2) They protect host plants against foreign proteases produced by pathogens and herbivores (reviewed in (Fluhr et al., 2012; Jamal et al., 2013; Kehr, 2006; Ryan, 1989; Zhu-Salzman & Zeng, 2015)).

Based on the specific active binding sites in the protein sequences, PINs can be classified into four main families: cysteine protease inhibitors, metalloid protease inhibitors, aspartic protease inhibitors, and serine protease inhibitors (Laskowski & Kato, 1980). Among the PIN families found in plants, serine proteinase inhibitors (SerPINs) have received the most attention, due to their functions in defense against insect herbivories and their dissimilarity from other PINs (Jamal et al., 2013; Ryan, 1989). Based on differences in their sequences and modes of action, SerPINs can be further divided into thirteen gene families, including the Potato PIN I/II families, Soybean trypsin inhibitor (Kunitz) family, Bowman-Birk family, barley trypsin inhibitor family, and squash inhibitor family (Birk, 2003; Ryan, 1989).

Since the initial demonstration of PIN activity in plant resistance against Colorado potato beetles (Green & Ryan, 1972), numerous studies have shown that PINs protect host plants against a wide range of insect species, including bruchid beetles (*Callosobruchus maculatus*) (Gatehouse & Boulter, 1983) and several lepidopteran species: tobacco budworm (*Heliothis virescens*) (Hilder et al., 1987), tobacco hornworm (*Manduca sexta*) <u>(Johnson et al., 1989)</u>, cotton leafworm (*Spodoptera littoralis*) (Tamayo et al., 2000), cotton bollworm (*Helicoverpa armigera*) (Udamale et al., 2013), fall armyworm (*S. frugiperda*), and beet armyworm (*S. exigua*) (Smigocki et al., 2013). While the significance of protein digestion for sap-feeding insects is still unclear, PINs are found in phloem sap of different plant species (Kehr, 2006), and PINs are effective in protecting plants against phloem-feeding insects, including several aphid species <u>(Bhatia et al., 2012; Casaretto & Corcuera, 1998; Pyati et al., 2011; Rahbé et al., 2003; Rausch e</u>t a<u>l., 2016; Tran et al., 1997)</u>.

While many PINs are effective in protecting plants against specific herbivores, others are not as effective due to their protease substrate specificity and the development of insect resistance. For example, aphid feeding induced PINs in barley (Casaretto & Corcuera, 1998) but not in tomato (Stout et al., 1998). Also, while potato PIN-I and PIN-II increased mortality of cereal aphids (*Diuraphis noxia*, *Schizaphis graminum*, and *Rhoalosiphum padi*), a lima bean PIN showed limited effects on any of these three cereal aphids (Tran et al., 1997). In the course of the arms race between insects and plants, insects can also develop resistance mechanisms against plant PINs (reviewed in (Jamal et al., 2013; Zhu-Salzman & Zeng, 2015)). The currently identified resistance mechanisms include overexpression of the PIN-sensitive protease and/or expression of other proteases (Ahn et al., 2004; De Leo et al., 1998), expression of PIN-insensitive proteases to avoid PIN digestion (Bolter & Jongsma, 1995; Oppert et al., 2005; Spit et al., 2012), and expression of proteases that digest the ingested PINs (Girard et al., 1998; Yang et al., 2009). Insects can combine multiple strategies to combat plant PIN defenses <u>(Zhu-Salzman et</u> <u>al., 2003)</u>. It is therefore a challenge for agronomists to identify potent PIN candidates that limit insect compensatory effects.

Although it is yet to be determined how insects sense the dietary PINs and respond accordingly, the identification and expression of non-host PINs is an effective strategy against the compensatory counter-defenses of resistant insects (Smigocki et al., 2013). For instance, transgenic tobacco (*Nicotiana tabacum*) plants expressing tomato and potato PINs reduce the growth of *M. sexta* (Johnson et al., 1989). Similarly, transgenic expression of a serine PIN from the American black nightshade, *Solanum americanum*, in *N. tabacum* enhanced resistance to *H. armigera* and the tobacco cutworm, *S. litura* (Luo et al., 2009). Expression of a maize PIN in rice plants increased resistance to the striped stem borer, *Chilo suppressalis*, by reducing insect growth (Vila et al., 2005). Although, transgenically expressed non-host PINs are often effective against target insects, some do not produce any effects or even have positive effects. For example, expressing the mustard trypsin PIN-II in *N. tabacum* did not enhance resistance to the *S. littoralis*, but instead promoted development and increased insect weight (De Leo et al., 1998). Those results suggest that the selection of effective non-host PINs requires further research effort. Advanced genome sequencing technology has generated growing numbers of plant genomes, which provide an enormous resource for identifying and expanding the useful repertoire of plant PINs. Furthermore, rapidly developing transgenic methods, *e.g.* CRISPR/Cas9 (Ellison et al., 2020), facilitate the functional characterization of candidate PINs for targeting specific insects.

Here we describe the repertoire of *M. persicae* proteases and show that, in comparison to whole aphids, a specific serine protease is highly expressed in the gut. In a VIGS screen of the non-preferred host plant, *N. benthamiana*, we observed that aphid survivorship was significantly increased by knockdown of a proteinase inhibitor PIN-II gene. To identify *N. benthamiana* PIN-IIs for the development of aphid control strategies, we annotated three *SerPIN-II* genes in the sequenced *N. benthamiana* genome (Bombarely et al., 2012; Kurotani et al., 2023) and characterized their role in plant defense against aphids using VIGS with TRV (Feng & Jander, 2022), CRISPR/Cas9 gene-editing (Ellison et al., 2020), and turnip mosaic virus (TuMV)-mediated overexpression (Casteel et al., 2014, 2015).

## Materials and Methods

### Aphid and plant cultures

The tobacco adapted green peach aphid strain, *Myzus persicae* (USDA-Red) (Ramsey et al., 2007, 2014) was maintained on *Nicotiana tabacum* at 23 °C with a 16:8 h light:dark photoperiod. Wildtype, *Cas9*-expressing (Baltes et al., 2015), and mutant *N. benthamiana* plants were maintained in a growth chamber with a 23 °C and a 16:8 h light:dark photoperiod. All plants were grown in Cornell Mix (by weight 56% peat moss, 35% vermiculite, 4% lime, 4% Osmocote slow-release fertilizer [Scotts, Marysville, OH], and 1% Unimix [Scotts]) in a Conviron (Winnipeg, Canada) growth chamber.

### Initial screening of proteinase inhibitor in aphid resistance with VIGS

Three-week-old *N. benthamiana* plants were inoculated with *Agrobacterium tumefaciens* strain GV3101 carrying tobacco rattle virus (TRV) plasmids. Aphids from *N. tabacum* were transferred and caged to *Agrobacterium*-infiltrated *N. benthamiana* plants when the plants were about seven weeks old. The cages contained five aphids with two cages per plant, though some plants held only one cage of five aphids. The aphids were of varied ages, with a consistent size. On the seventh day after placing the aphids on the plants, we counted the number of aphids to measure survivorship. Two to four cages of aphids were harvested for each VIGS-silenced genotype.

### *Identification of proteinase inhibitors from* N. benthamiana *genome*

To annotate the proteinase inhibitors, the previously identified *N. benthamiana PIN-II* (GeneBank ID: DQ158182.1) was used to BLAST against the *N. benthamiana* genome on Sol Genomic Network (Bombarely et al., 2012) and the newly released version (Kurotani et al., 2023), from which two PIN-IIs were identified, Niben101Scf09186g00005.1 and Niben101Scf09186g00006.1. In addition, the potato (*Solanum tuberosum*) *PIN-II* (GenBank ID: AIT42246.1) was used to BLAST against the *N. benthamiana* genome, and identified another PIN-II, Niben101Scf07757g00001.1. All sequences were reciprocally BLASTed against NCBI to confirm their identities.

### Proteinase inhibitors phylogeny

To infer the functions of *N. benthamiana* PINs, we used the maximum likelihood method to construct a protein phylogenetic tree of previously annotated Solanaceae PINs. We collected all available protein sequences of Solanaceae PIN families (including proteinase inhibitor I, Trypsin I, Multicystatin, Kunitz cysteine PINs, Aspartic PINs, and PIN-IIs) from the MEROPS the peptidase database Release v12.4 (https://www.ebi.ac.uk/merops/inhibitors/) (Rawlings et al., 2018). Before constructing the phylogenetic tree, we removed all sequences without assigned functional groups, as well as duplicated and within-species homologous (identity > 90%). The retained sequences were aligned in ClustalW (Thompson et al., 1994). Then the alignment was improved by removing poorly aligned regions with a gap threshold 0.25 using TrimAl v1.4 (Capella-Gutiérrez et al., 2009). Finally, a unrooted maximum likelihood tree was built with a bootstrap of 1000 in RAxML v8.2.12 (Stamatakis, 2014). To construct a phylogenetic tree of Solanaceae PIN-IIs, we followed the same protocol, with additional available PIN-II sequences that were obtained from NCBI (https://www.ncbi.nlm.nih.gov). The phylogenetic trees were visualized and presented using FigTree v1.4.4 (http://tree.bio.ed.ac.uk/software/figtree/).

### *CRISPR/Cas9 mutagenesis of* N. benthamiana

To generate stable PIN mutants, three single-guide RNAs (sgRNA) were designed for each PIN-II gene using the CRISPRdirect (https://crispr.dbcls.jp/). SgRNAs with no potential off-target sites with 20-mer matches and >45% GC content were selected. The gRNAs were cloned into a modified TRV vector following the TRV gRNA Golden Gate Assembly protocol (Ellison et al., 2020). Briefly, the gRNA_forward_primer (acacctgcaacgAAACxxxxxxxxxxxxxxxxxxxxGTTTTAGAGCTAGAAA, with x denoting the guide sequence) and the final_reverse primer (tcacctgctagtCACTctaaagtcttcttcctccgc) were synthesized by IDT (Integrated DNA Technologies, IA, USA). PCR amplifications were set up using the Q5 protocol (New England BioLabs, MA, USA), with the previously described plasmid (pEE392, (Ellison et al., 2020)) as a template. The PCR amplifications were treated with *Dpn*I (New England BioLabs, MA, USA) to remove the template plasmid. The purified PCR amplicons were then used to set up a Golden Gate reaction (Ellison et al., 2020) to clone this gRNA into the destination vector pEE083. The Golden Gate reaction was as follows: 50 ng pEE083 plasmid, 0.5 μl of PCR amplicon, 0.4 μl *Aar*I oligonucleotide, 0.5 μl *Aar*I enzyme, 1 μl T4 DNA ligase, 2 μl 10 x T4 DNA ligase buffer, and distilled H_2_O up to 20 μl. The PCR program was: 10 cycles of 37 °C for 5 min and 16 °C for 10 min, followed by 37 °C for 5 min, 80 °C for 5 min, and 4 °C hold. The Golden Gate assembly reaction was then transformed into *E. coli* and positive colonies were selected using kanamycin and confirmed by Sanger sequencing using the RN321 and RN322 primers (RN321: gggatttaaggacgtgaactc, and RN322: gtagtttaatgtcttcgggac). Plasmids with gRNAs inserted were further transformed into *Agrobacterium* strain GV3101, along with a TRV1 plasmid (Senthil-Kumar & Mysore, 2014), and gRNAs were infiltrated into *cas9-*expressing *N. benthamiana* (Ellison et al., 2020) to induce the mutagenesis. Seeds from the infiltrated *N. benthamiana* were planted and screened for homozygous mutants of proteinase inhibitors using PCR followed by confirmation with Sanger sequencing.

### *Overexpression of proteinase inhibitor II in* N. benthamiana

PINs were overexpressed using a modified Turnip mosaic virus (TuMV) vector (Casteel et al., 2014, 2015). The full length of SerPIN-II1/2 coding sequences with the cloning elements were synthesized with by GeneWiZ (https://web.genewiz.com/) after a codon optimization, due to the repeated fragments within the sequences and the high sequence similarity between the two *SerPIN-II* genes. The full-length *SerPIN-II3* coding sequences were amplified from *N. benthamiana* cDNA. The primers were designed to ensure an in-frame coding sequence and removal of the stop codon, as well as adding specific enzyme cutting sites (*Nhe*I and *Nco*I) to facilitate the cloning of the amplicon to the empty TuMV vector. The synthesized *SerPIN-II1/2*, cloned *SerPIN-II3*, and the TuMV empty vector were first subjected to double digestion with *Nhe*I and *Nco*I at 37 °C for 20 min, and then the enzyme activities were inactivated at 80 °C for 20 min. Following purification of the digestion products, the digested *SerPIN-II*s were ligated with the digested TuMV empty vector using T4 DNA ligase at room temperature for 15 min and inactivated at 65 °C for 20 min. The successfully ligated constructs were transformed to *E. coli*, with kanamycin selection, and then bacterial colonies were amplified and screened for positive colonies using Sanger sequencing. The positive constructs were transformed into *Agrobacterium* GV3101, and infiltrated into ∼4-week-old *N. benthamiana* for overexpression. A TuMV that expresses GFP (TuMV-*GFP*) was constructed and infiltrated into plants in parallel as control. After ∼10 days, the over-expression of *SerPIN-IIs* was confirmed using qPCR, and GFP expression was checked under a fluorescence microscope (Leica M205 stereomicroscope, in the Plant Cell Imaging Center at the Boyce Thompson Institute).

### Aphid bioassays

Aphid bioassays were performed as previously described (Feng et al., 2021). For *serpin-II3* mutant plants, 4 to 5-week-old plants were used for the bioassays, with *Cas9*-expressing plants used as controls. For the overexpression plants, ∼4-week-old *N. benthamiana* were infiltrated, and 10 days after the infiltration, each plant was confirmed for *SerPIN-II3* overexpression and then used for bioassays. In each experiment, 8 plants were used for each mutant or construct, and each experiment was repeated two to four times.

For each mutant or construct, two cages were set up per plant for the aphid bioassay. Ten to 15 adult aphids were placed in each cage and allowed to produce nymphs for ∼12 hours. Then, all adults were removed and 25 nymphs were left in each cage to monitor aphid survival every day for 5 days. At the end of the survival monitoring, five adult aphids were left in each cage to monitor aphid reproduction for a week. The rest of the aphids were collected and imaged using a Leica M205 stereomicroscope (Plant Cell Imaging Center, Boyce Thompson Institute), and aphid sizes were assessed by measuring the two-dimensional surface area of each aphid using ImageJ (Schneider et al., 2012). After images were taken, aphids were collected with 20 aphids per plants as one sample, weighed, frozen in liquid nitrogen, and stored at -80 °C for protein content measurements.

### Aphid protein content measurement using Bradford assays

For Bradford assays, the aphid samples were first crushed using a tissue homogenizer (1600 MiniG, SPEX^@^SamplePrep, NJ, USA), and then 20 times (volume in μl/weight in mg) of lysis buffer (100 mm KH_2_PO_4_, 1 mm dithiothreitol, and 1 mm ethylenediaminetetraacetic acid, pH 8.0) were added into each sample and mixed well by vortexing. Mixed samples were immediately centrifuged at 200 x g for 5 mins and 20 μl of the supernatant was transferred into 1 ml Bradford reagent (Bio-Rad, CA, USA) in 1.5 ml tubes. The samples were incubated at room temperature for 20 minutes and protein content was measured using a spectrophotometer (Model: Cary 60 UV-Vis, Agilent Technologies, CA, USA) at 595 nm. In parallel, a serial dilution of bovine serum albumin (BSA) with known concentrations was used to generate a standard curve for the quantification of the aphid samples. Lastly, the aphid protein content (mg) was normalized to the total amount of aphid sample (mg) and the number of aphids to allow comparisons across different treatments.

### Aphid gut protease expression

To identify aphid gut proteases that are potentially targeted by plant PINs, we annotated protease genes in the newly sequenced *M. persicae* (strain USDA-Red) genome (Feng et al., 2023) and profiled their expression in whole aphids and aphid guts using previously published transcriptomic data from three aphid lineages (USDA-Red, G002, G006) collected from cabbage (*Brassica oleracea*), a preferred host plant (Feng et al., 2018). Sequences of aphid proteases were obtained from NCBI (https://www.ncbi.nlm.nih.gov) and BLASTed against the protein models of the *M. persicae* USDA-Red genome (Feng et al., 2023). All identified *M. persicae* proteases were reciprocally BLASTed against the NCBI protein database to confirm their identities, and sequences with ambiguous annotations were removed from further analyses. To extract expression information for the *M. persicae* protease genes, the corresponding DNA sequences were blasted against the transcripts that were *de novo* assembled from the transcriptome data of three *M. persicae* lineages (Feng et al., 2018). Finally, using methods described previously (Feng et al., 2018), *PIN* gene expression levels were quantified, including the log_2_FC (fold change), p-values in the differential expression analyses, and the normalized count per million reads (CPMs) in each sample.

### Statistical analyses

All statistical comparisons were performed using R studio v1.3.959. For the aphid survival, size, reproduction, and protein content, data were combined from each repeated experiment for generating figures and statistical analyses. We tested for differences using a generalized linear model with a fixed factor of treatments and experiments as a block effect, followed by a Dunnett’s post hoc test to compare each treatment (if multiple groups) to the corresponding control group.

## Results

### Identification of gut-expressed aphid proteases

Although plant-produced PINs have been implicated in resistance to aphid feeding <u>(Bhatia et al.,</u> <u>2012; Casaretto & Corcuera, 1998; Pyati et al., 2011; Rahbé et al., 2003; Rausch et al., 2016;</u> <u>Tran et al., 1997)</u>, we are not aware of a comprehensive analysis to identify proteases that are expressed specifically in aphid guts. Using the recently sequenced *M. persicae* strain USDA-Red genome (Feng et al., 2023) and previous published transcriptome data from three *M. persicae* lineages (USDA-Red, G006 and G002) (Feng et al., 2018), we identified 82 protease genes. For 67 of these protease genes, expression was reliably detected in whole aphids and/or dissected gut tissue (with a cpm > 2 in over 3 samples) (Figure 1). Eight protease genes were expressed at a significantly higher level in the aphid guts than in whole insect tissue (Figure 1A). Two of those gut-expressed genes (MPE009914 and MPE002313, highlighted in red in Figure 1B) were annotated as serine proteases. In addition, the serine protease MPE009914 had the highest gut expression level among all 67 proteases in all three tested *M. persicae* lineages. This expression level is even higher than that of the previously characterized and abundant cathepsin MPE001251 (Rahbé et al., 2003; Rauf et al., 2019), which has the overall second-highest expression level (highlighted in red in Figure 1B). In addition to the abundant cathepsin L and gut-specific serine protease, we identified several other gut-enriched proteases, including another serine protease (MPE002313), calpain-A/C (MPE012440/MPE023509), an aspartic protease (MPE001815), a cysteine protease (MPE011270), an ion protease (MPE000063), and a Clp protease (MPE017073).

**Figure 1.**
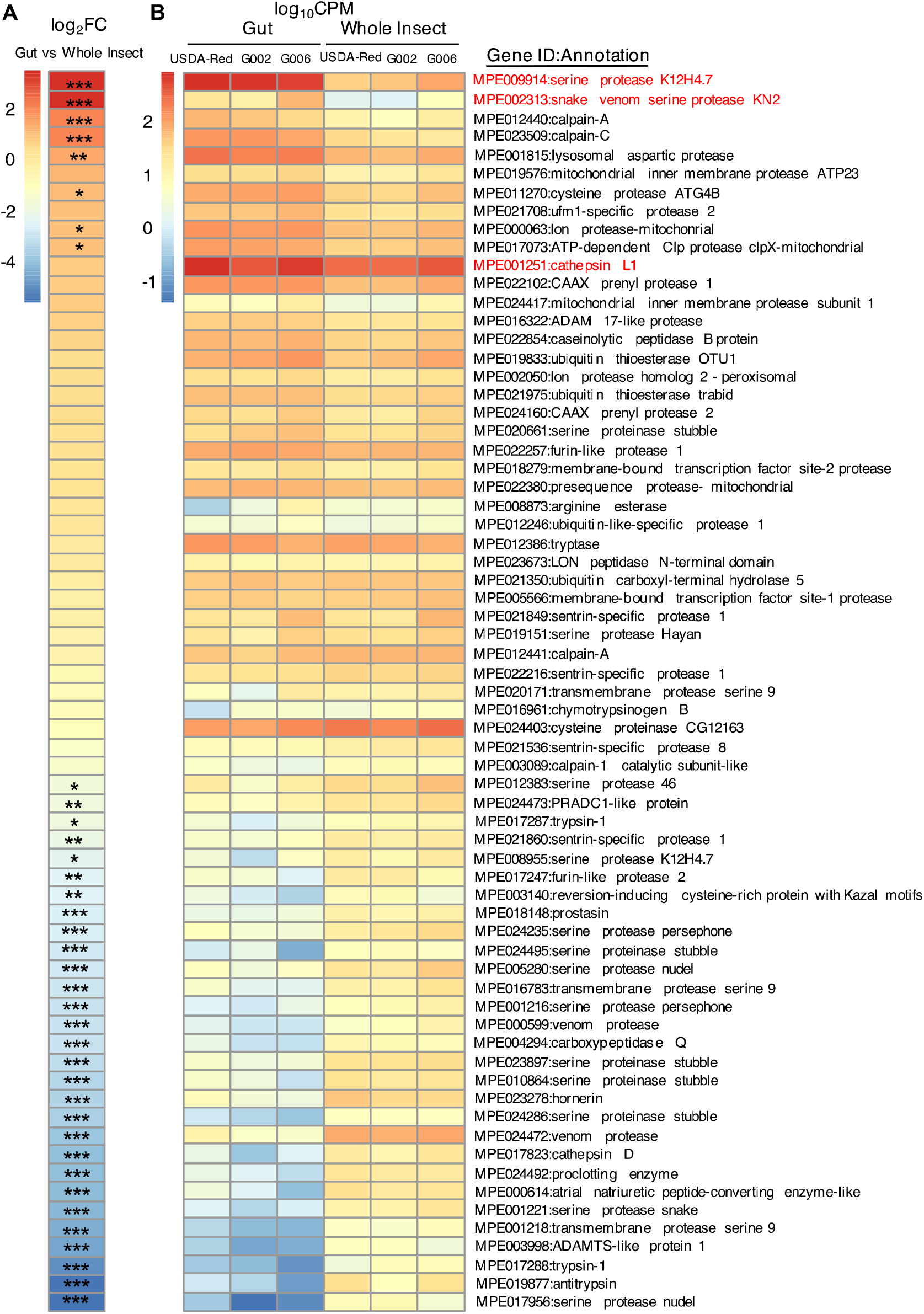
Annotation of aphid proteinases and their relative expression in the gut. A list of 67 aphid proteases were annotated in the genomes of three *M. persicae* lineages (USDA-Red, G002, and G006). (A) Fold change (Log_2_FC) of aphid proteinase gene expression, comparing expression levels in the gut and the whole insects. Proteases are ordered from highest to lowest relative expression levels, and significant differences in the expression levels between aphid guts and whole aphids are indicated by: *\* p* < 0.05, ** *p* < 0.01, and *** *p* < 0.001. (B) Read counts for each gene were normalized to Log_10_CPM (count per million reads) to show the relative abundances of aphid protease genes across tissue types and aphid clones. Red highlighted text indicates two serine proteinases that are significantly upregulated in aphid guts comparing to the whole insects, and one cathepsin L1 gene that is enriched in aphid gut (ranked No. 2 in log_10_CPM across all proteinases in the gut), but not expressed at a significantly higher level in the guts compared to the whole insect samples. Cathepsin activity, but not serine protease activity, has been previously reported in aphid guts (Rahbé et al., 2003; Rauf et al., 2019). Gene identifiers in this figure are from the newly sequenced *M. persicae* genome (Feng et al., 2023).

### PIN-II affects aphid survival

In a separate approach, we conducted aphid experiments using a VIGS library that was initially developed for screening pathogen resistance genes in tomato and *N. benthamiana* (Pozo et al., 2004; Senthil-Kumar et al., 2018). Among 110 defense-related genes that were expression-silenced in these experiments, a VIGS construct based on the tomato *PIN-II* gene (SHR58 in Figure S1) elicited the greatest increase in *M. persicae* survival. Compared to the empty vector control there was an almost three-fold increase (p < 0.01, Figure 3A) in the number of aphids on plants with the tomato *PIN-II* (SHR58) VIGS construct <u>(Pozo et al., 2004)</u>. Although we did not quantify *N. benthamiana* gene expression levels after the VIGS treatment, we considered this an indication that *N. benthamiana* PINs contribute to aphid defense and merit further investigation.

### *Identification of type II serine proteinase inhibitors (SerPIN-IIs) in* N. benthamiana

Using reciprocal blast, we identified three members of the proteinase inhibitor II family in *N. benthamiana.* Since a phylogenetic tree of the major protease inhibitor gene families (Figure S2) classified all three of our candidate proteins in the PIN-II group, we further classified our candidate PINs with a more detailed phylogenetic tree of annotated Solanaceae serine protease inhibitors (Figure 2). The identified *N. benthamiana* gene (Niben101Scf09186g00005.1, Niben101Scf09186g00006.1, and Niben101Scf07757g00001.1 (marked with red text in Figure 2) all have potato type II functional domains (Figure S2). We therefore named these genes *SerPIN-II1*, *SerPIN-II2*, and *SerPIN-II3*, respectively. Further phylogenetic analyses revealed that *SerPIN-II1* (Niben101Scf09186g00005.1) and *SerPIN-II2* (Niben101Scf09186g00006.1) were most closely related to trypsin proteinase inhibitor II sequences, whereas *SerPIN-II3* (Niben101Scf07757g00001.1) was clustered with PIN-IIs that have no specific functional annotations (Figure 2).

**Figure 2.**
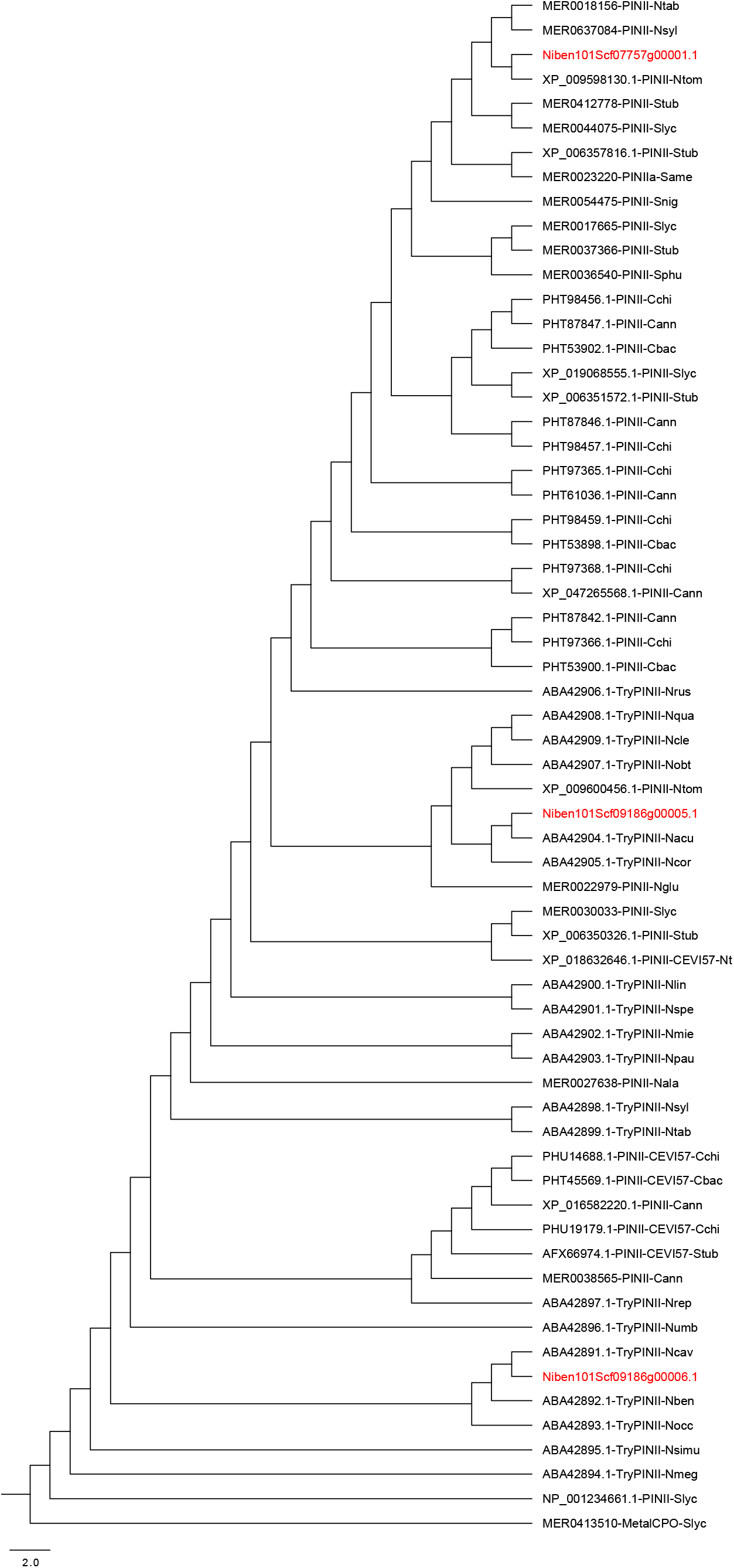
Phylogeny of type II proteinase inhibitors (PIN-II) from the Solanaceae family. A rooted maximum likelihood tree was built using RAxML with 1000 bootstraps with MER0413510-MetalCPO-Slyc as an outgroup. To help visualize the branches, the phylogenetic tress was transformed into cladogram. The three *N. benthamiana* serine PIN-IIs identified in this study are highlighted in red. All PIN-II proteins in the tree are within the major serine PIN-II family, containing one or multiple potato PIN-II functional domains. Both Niben101Scf09186g0005.1 and Niben101Scf09186g0006.1 were grouped with trypsin-type PIN-IIs, whereas the Niben101Scf07757g00001.1 was grouped among PIN-II genes without further functional annotations. The naming of each protein has three parts, protein IDs from either the MEROPS database (MER#######) or NCBI, followed by the PIN-II name (*e.g.,* TryPINII), and the plant species. The Solanaceae plant species collected included, Cann: *Capsicum annuum*; Cbac: *C. baccatum*; Cchi: *C. chinensee*; Nacu: *Nicotiana acuminata*; Nala: *N. alata*; Nben: *N. benthamiana*; Ncav: *N. cavicola*; Ncle: *N. clevelandii*; Ncor: *N. corymbosa*; Ngla: *N. glauca*; Nglu: *N. glutinosa*; Nlan: *N. langsdorffii*; Nlin: *N. linearis*; Nmeg: *N. megalosiphon*; Nmie: *N. miersii*; Nobt: *N. obtusifolia*; Nocc: *N. occidentalis*; Npau: *N. pauciflora*; Nqua: *N. quadrivalvis*; Nrep: *N. repanda*; Nspe: *N. spegazzinii*; Ntab: *N. tabacum*; Ntom: *N. tomentosa*; Nrus: *N. rustica*; Nsim: *N. simulans*; Nsyl: *N. sylvestris*; Numb: *N. umbratica*; Same: *S. americanum*; Sber: *S. berthaultii*; Sbre: *S. brevidens*; Slyc: *S. lycopersicum*; Snig: *S. nigrum*; Sphu: *S. phureja*; Stub: *Solanum tuberosum*.

### *SerPIN-II1 and SerPIN-II2 confer* N. benthamiana *resistance to aphid feeding*

In the initial VIGS screening experiments (Figures S1 and 3A), the tomato PIN-II sequence was identical to *SerPIN-II2*, and was also predicted silence expression of *SerPIN-II1*, which has a high level of similarity and multiple regions with 19 base pairs of identity (Figure S3). However, due to the repeated sequences in the coding region, as well as the high similarity between *SerPIN-II1* and *SerPIN-II2* (Figure S3), we were not able to design gene-specific primers for qPCR experiments to confirm whether there was knockdown of both *SerPIN-II1* and *SerPIN-II2*.

We attempted to create *SerPIN-II1* and *SerPIN-II2* mutants using CRISPR/Cas9. However, again due to the repeated sequences in the coding region and the high similarity between the *SerPIN-II1* and *SerPIN-II2* (Figure S2), we were not able to confirm the mutations by Sanger sequencing or qRT-PCR. Nevertheless, our VIGS experiments (Figure 3A) demonstrated that *SerPIN-II1* and/or *SerPIN-II2* knockdown promotes against aphid growth and reproduction on *N. benthamiana*.

**Figure 3.**
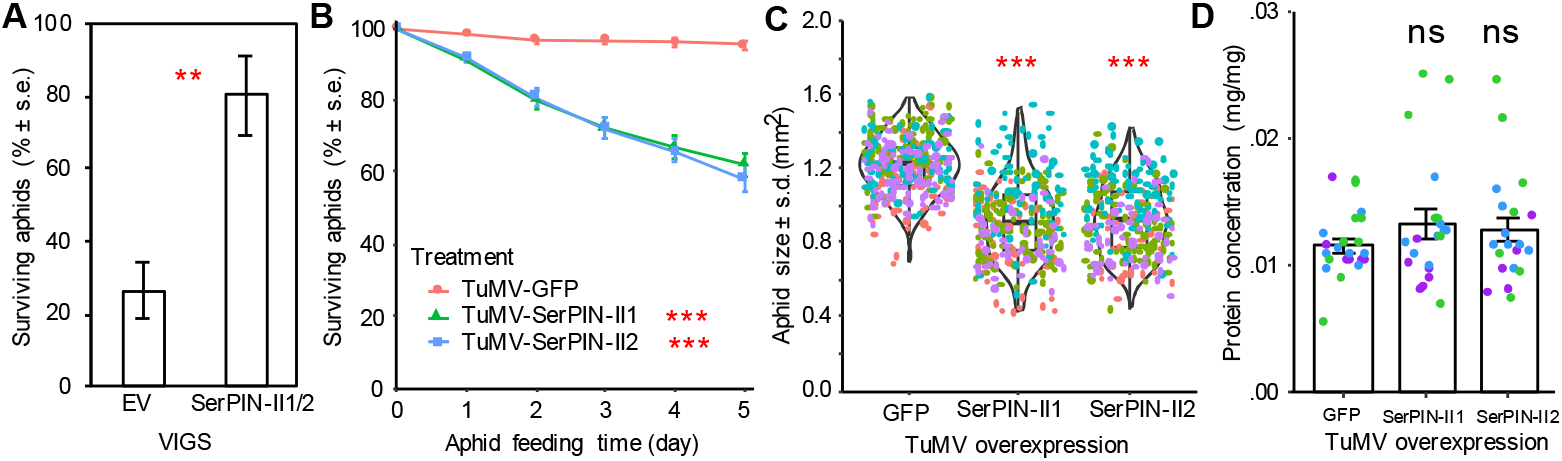
SerPIN-II1 and SerPIN-II2 confer *Nicotiana benthamiana* resistance to aphid feeding. (A) Aphid survival on empty vector control (EV) *N. benthamiana* and after tobacco rattle virus (TRV) virus induced gene silencing (VIGS) targeting *SerPIN-II1* and *SerPIN-II2*. Mean +/- s.e. of n = 14 (EV) and n = 8 (*PIN-II* VIGS), **P < 0.01, *t*-test. (B) Aphid survival after feeding on *N. benthamiana* with *SerPIN-II1* and *SerPIN-II2* overexpressed using a turnip mosaic virus (TuMV) vector. GFP overexpression was used as the negative control. Aphid bioassays were performed four times. Data shown are mean +/- s.e. of n = 56. Significant differences are indicated for the 5-day time point. (C) *N. benthamiana SerPIN-II1* and *SerPIN-II2* overexpression effects on aphid growth in four independent experiments (different colored dots are data from independent experiments; n = 300-350). (D) Aphid protein content was measured using Bradford assays in three of the four experiments. Mean +/- s.e. of n = 32. Significance was tested using a generalized linear model with a fixed factor of treatment and experiment as a block effect, ns = not significant, *\* p* < 0.05, ** *p* < 0.01, and *** *p* < 0.001.

When we overexpressed *SerPIN-II1* and *SerPIN-II2* in *N. benthamiana* using TuMV, aphid survival on the overexpression plants was significantly reduced compared to control TuMV-*GFP* plants (Figure 3B). In addition, aphid body size was significantly smaller on the *SerPIN-II1* and *SerPIN-II2* overexpression plants (Figure 3C). Due to the retarded growth, most of the aphids on the *SerPIN-II1* and *SerPIN-II2* overexpression plants were in juvenile stages, which did not allow us to perform aphid reproduction comparisons to the mature aphids on TuMV-*GFP* control plants. Furthermore, by the end of the bioassays, we did not observe significant differences in aphid protein concentrations after they had fed on *SerPIN-II1* and *SerPIN-II2* overexpressing plants and TuMV-*GFP* control plants (Figure 3D).

### *Aphid survival and growth on* SerPIN-II3 *mutant* N. benthamiana

We infected Cas9-expressing *N. benthamiana* with TRV carrying two guide RNAs (gRNA) to generate four independent homozygous mutant plants in the subsequent generation: *serpin-II3-1*, *serpin-II3-2*, *serpin-II3-3*, and *serpin-II3-4* (Figure 4). *serpin-II3-1* has a three-nucleotide deletion at the gRNA2 cutting site; *serpin-II3-2* has a single nucleotide insertion at the gRNA2 cutting site, leading to a frameshift; *serpin-II3-3* has a 161-nucleotide deletion generated by gRNA2; and *serpin-II3-4* has a single-nucleotide insertion at gRNA3, leading to a frameshift.

**Figure 4.**
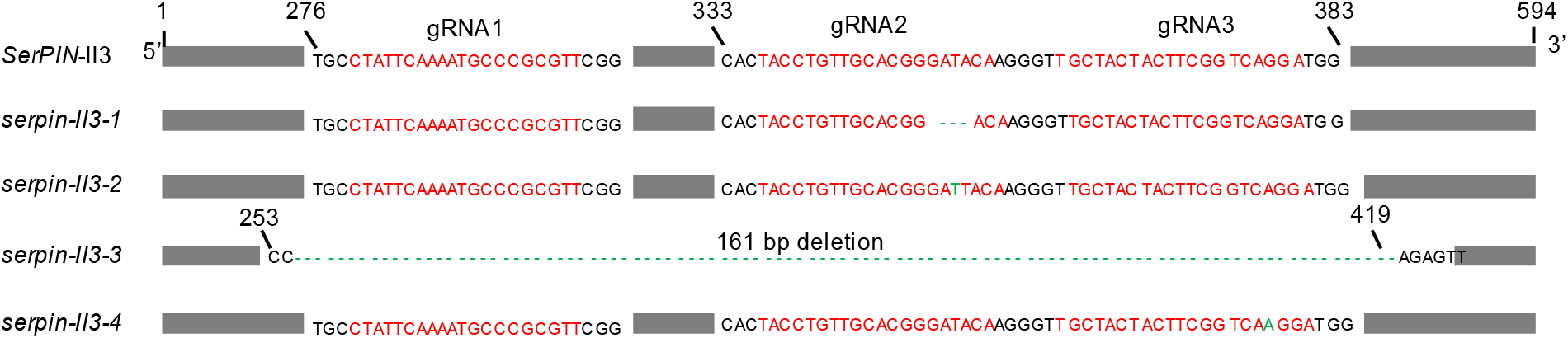
*Nicotiana benthamiana* serine type proteinase inhibitor 2 (*SerPIN-II3*) mutants were generated using CRISPR/Cas9. Three gRNAs (shown in red) were designed for mutagenesis, and gRNA2 and gRNA3 generated four independent mutations. Mutations (insertions and deletions) are highlighted in green. *serpin-II3-1* has a three nucleotides deletion at gRNA2, *serpin-II3-2* has a single nucleotide insertion at gRNA2, *serpin-II3-3* has a 161 nucleotide deletion generated by gRNA2, and *serpin-II3-4* has a single-nucleotide insertion at gRNA3.

To test the function of SerPIN-II3 in protecting *N. benthamiana* against insect pests, we performed bioassays with synchronized newborn *M. persicae* to monitor their survival, growth and reproduction. During the 5-day bioassay, we observed significant improvements in aphid survivorship when fed on *serpin-II3* mutants comparing to the *Cas9*-expressing control plants (*p* < 0.001, Figure 5A). After 5 days of feeding, surviving aphids on all four *serpin-II3* mutants were significantly larger than those on *Cas9*-expressing control plants (Figure 5B). At this point, five surviving aphids in each cage were allowed to reproduce for one week, but no significant differences were observed with aphids on *serpin-II3* mutants relative to control plants (Figure 5C). In the choice assays, we did not observe aphid preferences for the *serpin-II3* mutant *N. benthamiana* relative to the *Cas9*-expressing control plants (Figure S4).

**Figure 5.**
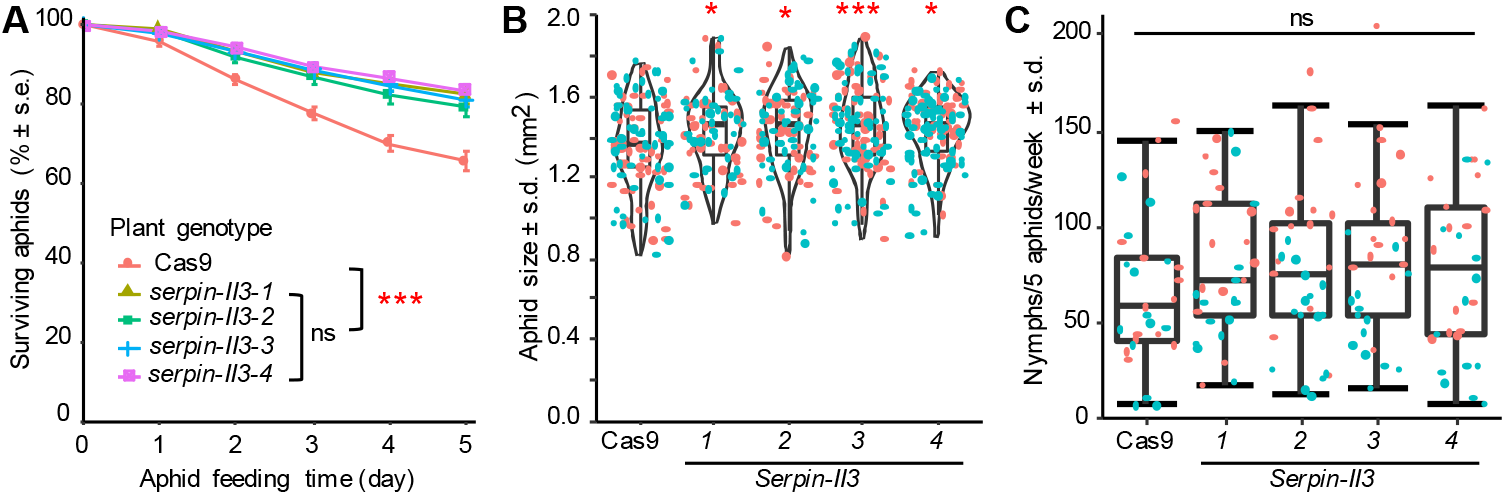
*SerPIN-II3* knockout mutations improve aphid survival and growth on *N. benthamiana*. Data presented in all panels were combined from two independent experiments (colored orange and cyan in panels B and C) (A) Aphid survival over 5 days, starting from newborn nymphs. Significant differences are indicated for the 5-day time point. Mean +/- s.e. of n = 15-16 per experiment. (B) Aphid growth, as measured by aphid size after 5 days of feeding on different genotypes of *N. benthamiana.* Each dot in the graph represents the size of a single aphid (n = 50-65 per experiment). (C) Aphid reproduction, as measured by the number of nymphs produced by 5 aphids (the surviving aphids from bioassays shown in panel A over one week, n = 15-16 per experiment). Significance was tested using a generalized linear model with a fixed factor of genotype and experiment as a block effect, followed by a Dunnett’s test for multiple comparisons. ns: not significant, * *p* < 0.05, ** *p* < 0.01, *** *p* < 0.001.

### *Aphid survival and growth on* SerPIN-II3 *overexpressing* N. benthamiana

To further confirm the function of SerPIN-II3 in protection against aphid feeding, we overexpressed *SerPIN-II3* using a turnip mosaic virus (TuMV) vector in wildtype *N. benthamiana* (*p* < 0.001, Figure 6A). Compared to *GFP*-overexpressing control plants (TuMV-*GFP*), aphid survival on the *SerPIN-II3* overexpression plants (TuMV-*SerPIN-II3*) was significantly decreased after five days (*p* < 0.001, Figure 6B). Aphids on TuMV-*SerPIN-II3* plants were significantly smaller than those on the TuMV-*GFP* plants (p < 0.001, Figure 6C). While most aphids on TuMV-*GFP* plants had reached maturity, most of the aphids on TuMV-*SerPIN-II3* plants were still in their juvenile stages. Therefore, there were not enough synchronized mature aphids to monitor aphid reproduction.

**Figure 6.**
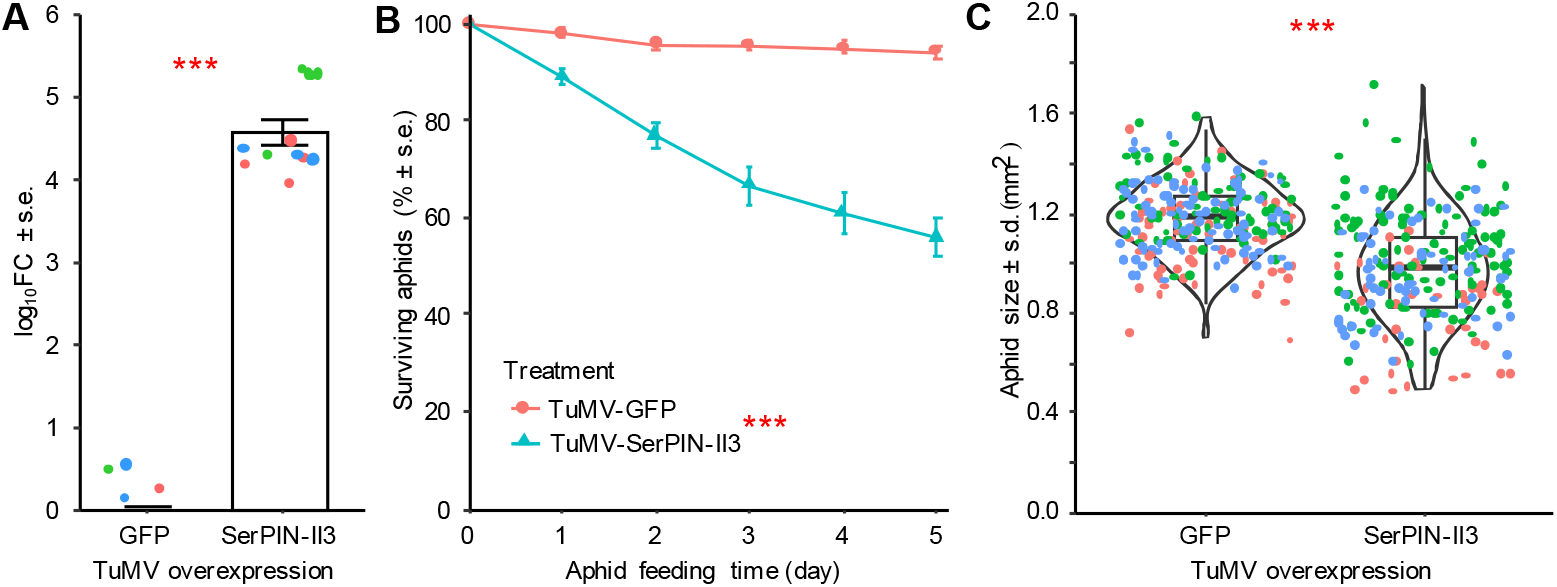
*SerPIN-II3* overexpression reduces aphid survival and growth. Data presented in all panels were combined from three independent experiments (colored red, green, and blue in panels A and C). (A) *SerPIN-II3* overexpression using a turnip mosaic virus vector (TuMV-*SerPIN-II3*). TuMV expressing green fluorescent protein (TuMV-GFP) was used as a negative control. Data represent log_10_FC of n = 5-6 plants for each experiment. (B) Aphid survival over 5 days starting from new-born nymphs. Significant differences are indicated for the 5-day time point, mean +/- s.e. of n = 10-12 cages per experiment (2-cages were used per plant). (C) Aphid size as a measure of aphid growth. Each dot represents the size of a single aphid (n = 70-100 aphids per experiment). Significance was tested using a generalized linear model with a fixed factor of genotype and experiment as a block effect, *\*\*\* p* < 0.001.

To exclude the possible effects of the native *SerPIN-II3* expression in the overexpression experiment, we further performed a rescue experiment, expressing *SerPIN-II3* on the four *serpin-II3* mutants (*e.g.*, *serpin-II3-1* infected with TuMV-*SerPIN-II3*). The SerPIN-II3 overexpression successfully complemented phenotype of *N. benthamiana* mutants (Figure 7). Aphid survival on the TuMV-*SerPIN-II3* plants was significantly reduced compared to TuMV-*GFP* control plants (*p* < 0.001), when overexpressed in the same *SerPIN-II3* mutant background (*p* < 0.001, Figure 7A, B, C, D).

**Figure 7.**
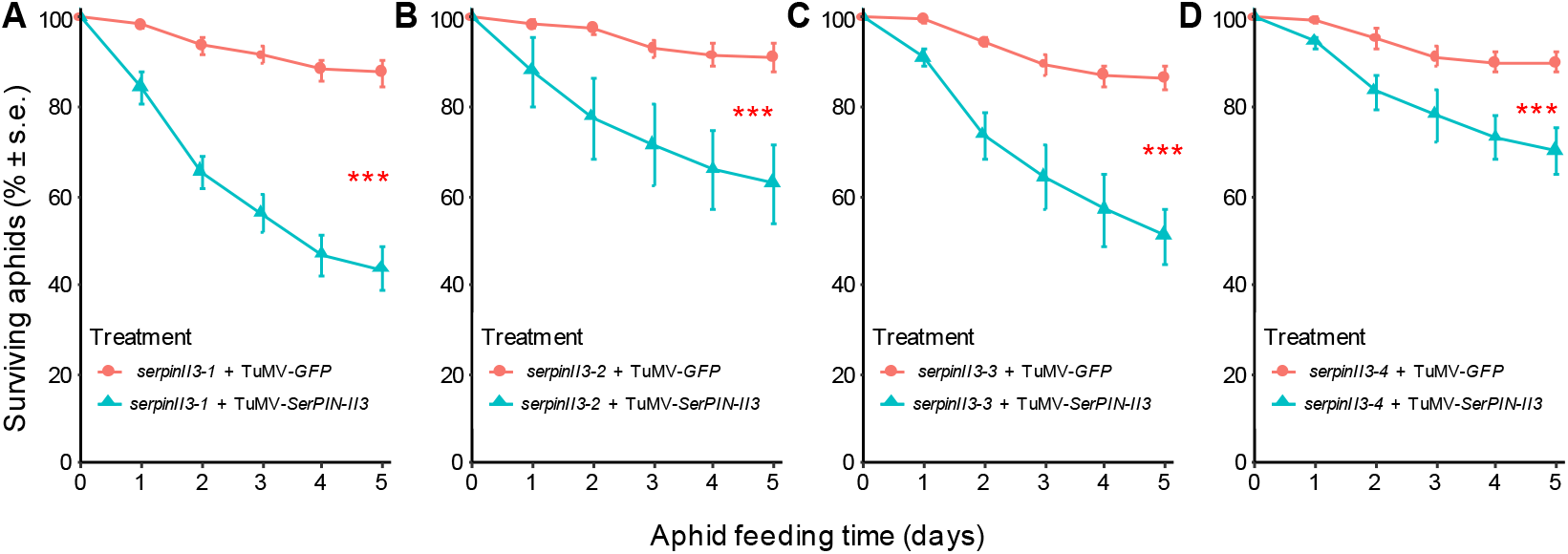
Resistance is restored by *SerPIN-II3* overexpression in *serpin-II3* mutant *N. benthamiana*. The SerPIN-II3 gene was overexpressed using a turnip mosaic virus (TuMV) vector in four *serpin-II3* mutant lines: (A) *serpin-II3-1;* (B) *serpin-II3-2;* (C) *serpin-II3-3;* (D) *serpin-II3-4.* n = 8 for each group. For the 5-day time point, significant differences are indicated, based on a generalized linear model with a fixed factor of treatment (equivalent to a two-sample *t*-test), *\*\*\* p* < 0.001.

### *SerPIN-II3 changes in* N. benthamiana *impact the aphid protein content*

As proteinase inhibitors function as inhibitors of proteolytic enzymes to digest proteins, we determined whether changes in the *SerPINII-3* expression level affect aphid protein content. After five days of feeding, aphids on the four *SerPIN-II3* mutants had significantly lower protein content than aphids fed on *Cas9*-expressing control plants (Figure 8A). Conversely, when we overexpressed *SerPIN-II3* in *N. benthamiana* (TuMV-*SerPIN-II3*), the aphid protein content was significantly increased, compared to aphids fed on TuMV-*GFP* control plants (Figure 8B).

**Figure 8.**
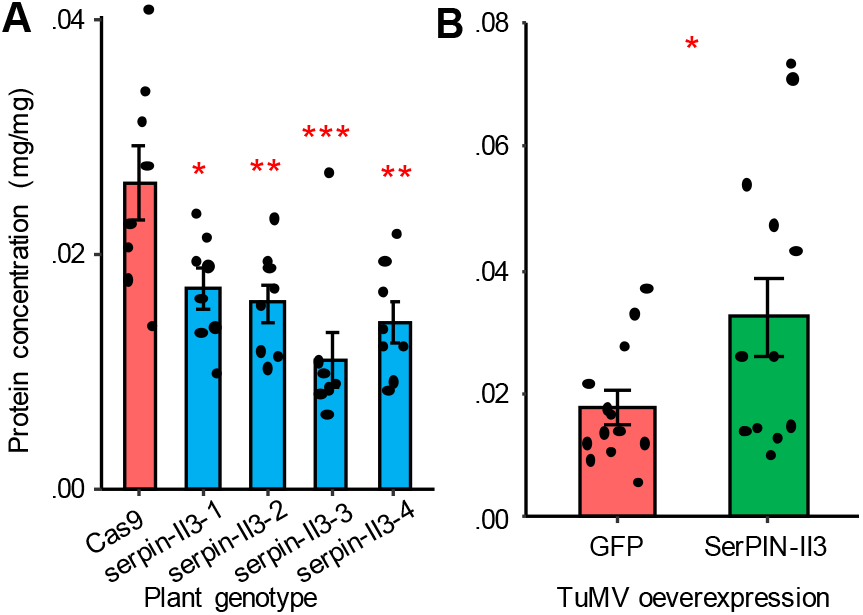
Aphid protein content measured after feeding on *N. benthamiana* plants. (A) Aphid protein content was significantly reduced after feeding on *serpin-II3* mutant *N. benthamiana* plants; a *Cas9*-expressing plant was used as the negative control, n = 8 (aphids collected from 8 different plants). Significance was tested using a generalized linear model with a fixed factor of plant genotype followed by a Dunnett’s test for multiple comparisons. * *p* < 0.05, ** *p* < 0.01, *** *p* < 0.001. (B) Aphid protein content significantly increased after feeding on *SerPIN-II3* overexpressing *N. benthamiana* plants; GFP overexpression was used as the negative control. The overexpression experiment was repeated twice, with n = 7-8 replicates per experiment. Significant differences were tested using generalized linear model with a fixed factor of treatment with experiment as a block effect, *\* p* < 0.05.

## Discussion

In this study, we identified three *N. benthamiana* genes encoding type II proteinase inhibitors (PIN-IIs) belonging to the main serine PIN family (*SerPIN-II1*, *SerPIN-II12*, and *SerPIN-II3*). Using VIGS, CRISPR/Cas9 gene-editing, and TuMV-mediated overexpression followed by aphid bioassays, we demonstrated that all three SerPIN-IIs from *N. benthamiana* can confer resistance to aphid feeding. As PINs are proteolytic enzymes that inhibit proteases, we further identified certain serine proteases were abundant in aphid gut as potential SerPIN-II targets, and showed that the protein content of aphids changed when aphids were fed on *PIN-II* mutants and/or *PIN-II* overexpressing plants.

While we observed significant effects on aphid growth after the knockdown (Figure 3A) and overexpression (Figure 3B, C) of *SerPIN-II1 and SerPIN-II2*, we were not able to confirm the mutations of our CRISPR/Cas9 mutants and/or confirm the downregulation/overexpression of our *SerPIN-II1 and SerPIN-II2*. The failure in our mutant confirmation was due to mixed signals during Sanger sequencing resulting from: 1) the repeat sequences in the gene (underlined in red in Figure S3) caused ambiguous sequences due to depletion, slippage, secondary structures, or 2) the high identity of *SerPIN-II1* and *SerPIN-II2* (Figure S3) could cause mutations in one gene could be blended with the other during gene amplification. *SerPIN-II1* and *SerPIN-II2* are located on the same scaffold in the *N. benthamiana* genome (Scf09186:179,955-182,349 for *SerPIN-II1* and Scf09186:129,137-149,078 for *SerPIN-II2*), and were possibly generated by tandem duplication. Although there are some unclear annotations in the genomic region (Figure S3), this sequencing problem could be solved using the more recent long-read genome sequencing technologies, such as PacBio and Oxford Nanopore Technologies (Marx, 2023) that would produce long reads to cover the full genomic regions of *SerPIN-II1* and *SerPIN-II2*.

We demonstrated that three *N. benthamiana SerPIN-IIs* reduce the survival and growth of *M. persicae*, and that these effects could be caused by the inhibition of specific aphid gut proteases. Our aphid gut proteases analyses identified several aphid proteases that are abundant and/or significantly upregulated in aphid gut comparing to the whole insects. Among those, a serine protease (MPE009914) is the most highly expressed gut-specific protease (Figure 1). However, previous publications suggested that cysteine proteases are the major type of digestive enzymes in aphid gut, as genes in the cathepsin families (*e.g.*, cathepsin B and L) have been identified as abundant gut specific cysteine proteases in the pea aphid, *Acythosiphon pisum* (Cristofoletti et al., 2003), the cotton aphid, *Aphis gossypii* (Deraison et al., 2004), and the cereal aphid, *Sitobion aveanae* (Pyati et al., 2011). Although a cathepsin L protease (MPE001251) was the second-most abundant in *M. persicae* gut tissue at the gene expression level, a similar expression level was detected in the whole insect (Figure 1). The high-level gene expression is consistent with observation that, when targeting cathepsin using cysteine proteinase inhibitors or RNAi, the survival, fecundity, and/or nutritional value of *M. persicae* are reduced (Rahbé et al., 2003; Rauf et al., 2019).

Prior to the current study, no gut-specific serine proteases were identified in aphids (Pyati et al., 2011). There has been an attempt to target a gut serine protease in *M. persicae* using RNA interference (Bhatia et al., 2012). However, the identification of the targeted gene was based on sequence homology to *A. pisum*, in which no gut specific serine protease was found (Pyati et al., 2011). Based on the published primer sequences, the serine protease target chosen by Bhatia et al. (2012) is MPE0023278 (hornerin). In our expression analysis, MPE0023278 is not abundantly expressed in the *M. persicae* gut (Figure 1), which may explain why there were no changes in *M. persicae* survival after gene expression knockdown (Bhatia et al., 2012).

Our identification of a gut-specific serine protease in *M. persicae*, coupled with previous indirect evidence, suggests that this difference in gut proteolytic activities could be species-specific. For example, overexpression of a *SerPIN* CI2c in barley, which is a non-preferred host for *M. persicae*, made plants more attractive for these aphids, perhaps through the suppression of other defenses (Losvik et al., 2018). Plant SerPINs affected the survival and reproduction of three cereal aphid species, *Diuraphis noxia*, *Schizaphis graminum*, and *Rhopalosiphum padi* (Tran et al., 1997). In *S. avenae*, although no specific serine proteases were identified, a recombinant wheat serine proteinase inhibitor, the subtilisin-chymotrypsin inhibitor (WSCI), led to more severe reductions in aphid survival and growth than a cysteine proteinase inhibitor, WCPI (Pyati et al., 2011). This phenomenon in *S. avenae* was partially explained by the fact that WSCI could also effectively inhibit insect cysteine proteases (Pyati et al., 2011). While the gut-specific serine protease of *M. persicae* (Figure 1) could be a major target of the *N. benthamiana* SerPIN-IIs, future research will be needed to further determine substrate specificity and spectrum of this aphid PIN.

As gut proteases are crucial for insects to digest proteins that ingested from their food sources, inhibition of these proteases will reduce nutrient uptake. For example, knockdown of gut cathepsin L using plant-mediated RNAi significantly reduced *M. persicae* body protein content, and more protein remained undigested and detected in the aphid honeydew (Rauf et al., 2019). Consistently, we also observed that *M. persicae* fed on *serpin-II3* mutants and *SerPIN-II3* overexpression plants have significantly changed protein contents, indicating effects on aphid proteolytic activities (Figure 8). However, the changes in protein content were opposite of what was observed with cathepsin L silencing. Knockout of *SerPIN-II3* reduced aphid protein content, and overexpression of *SerPIN-II3* increased aphid protein content (Figure 8). It is possible that the *serpin-II3* knockout mutants of *N. benthamiana* released plant-derived serine proteases in aphids, which could cause nutrient deficiencies by digesting aphid gut proteins. The fact that aphid sizes were larger on the *serpin-II3* mutant plants (Figure 5B) suggests that aphids ingested more phloem sap to compensate for the digestive activity of gut serine proteases, but the unit protein contents per body weight (mg/mg) for aphids was significantly reduced (Figure 8A). Conversely, when *SerPIN-II3* was overexpressed, the targeted proteases in aphids were digested, thereby the aphids reduced their further feeding of *SerPIN-II3* overexpressing plants to conserve endogenous proteins. Coupled with the fact that aphid sizes were smaller on the *SerPIN-II3* overexpression plants (Figure 6C), the unit protein content (mg/mg) for aphids was significantly increased (Figure 8B).

Although we observed significant protein content changes for *SerPIN-II3*, we did not observe significant protein content changes in aphids when overexpressing *SerPIN-II1 and SerPIN-II2*, even though the aphid sizes were significantly smaller when fed on *SerPIN-II1 and SerPIN-II2* overexpressing plants (Figure 3C, D). One explanation could be that the targets of *SerPIN-II1* and *SerPIN-II2* in the aphid gut are not involved in general proteolysis, other than inhibiting plant PINs. Secondly, aphids have been shown to secret salivary proteins, including proteases (e.g. M1/2 metalloprotease and aminopeptidase), effector proteins (e.g. Mp10 and Mp42), and detoxification enzymes (e.g. oxidoreductases and polyphenoloxidases, *etc*.), to suppress plant defense (reviewed in (Furch et al., 2015; van Bel & Will, 2016)). *SerPIN-II1* and *SerPIN-II2*, rather than inhibiting proteases in aphids, could inhibit the function of aphid-secreted salivary proteases in plants to promote defenses. Another explanation could be that the mode of action for *SerPIN-II1 and SerPIN-II2* is different from SerPIN-II3. Whereas SerPIN-II3 targets aphid proteases, *SerPIN-II1 and SerPIN-II2* may instead target plant endogenous proteases (Ryan, 1989) to prevent the proteolysis thereby limiting aphid’s nutrient access from feeding. Previous research has shown that some PINs participate in plant proteolysis, which can lead to programmed cell death and/or autophagy (Kim & Hwang, 2015; Lampl et al., 2013; Solomon et al., 1999), thereby limiting nutrient access at the infection sites for plant pathogens and/or insect herbivores. Although we did not observe any significant differences in whole-plant phenotypes between the *SerPIN-II1 and SerPIN-II2* overexpression plants and the control plants, differences in molecular and/or biochemical levels (*i.e*., programmed cell death or autophagy) at the aphid ingestion sites needs to be further evaluated. In addition, the induction of protease inhibitors by insect herbivores has been shown to be triggered by general plant defensive mechanisms, such as jasmonic acid signaling (Bolter & Jongsma, 1995; Xu et al., 1993), which is not directly related to proteolysis. All these possibilities need to be determined with further research.

In addition to the previously identified cathepsins B and L and the newly identified gut-specific serine proteases, we also identified several other gut-specific proteases, including a calpain-A/C, aspartic protease, an ion protease, and a Clp protease. While their exact functions still need to be characterized, this broad range of proteases could partially explain the plasticity of *M. persicae* as a generalist to be able to colonize a broad range of host plants (Mathers et al., 2017).

In addition to SerPIN-IIs, many PIN-Is from other plants have also been shown to provide defense against pathogens or insect herbivores (Martinez de Ilarduya et al., 2003; Pautot et al., 1991; Tran et al., 1997). Although some of PIN-Is were shown to be not as effective as PIN-IIs (Johnson et al., 1989), further research can be performed to identify PIN-Is and/or PINs of other types in *N. benthamiana*, and determine their functions in plant defenses. Regardless the lack of understanding of substrate specificities of PINs from biochemistry and enzymatic perspectives, overexpressing multiple PINs as a stack could be used to improve the efficiency of insect inhibition. For example, co-expression of two potato-type PINs (a *Nicotiana alata* PIN-II and a *Solanum tuberosum* PIN-Ia) in cotton yield better crop protection against the major lepidopteran cotton pests such as *Helicoverpa punctigera* and *H. armigera* (Dunse et al., 2010).

In addition to aphids, *N. benthamiana* is also a non-preferred host plant for several other tobacco-specialist and non-specialist insect pests, such as the corn earworm (*Helicoverpa zea*), tobacco budworm (*Heliothis virescens*), and cabbage looper (*Trichoplusia ni*) (Feng et al., 2021). Those insects can also be tested to determine the target-insect spectrum of non-host PINs in *N. benthamiana*.

Aphids are severe agricultural pests that cause damages to a wide range of crops. Researchers have strived to find novel technologies for aphid population control, but with limited success. By functionally characterizing three SerPIN-IIs from an *M. persicae* non-preferred host plant, *N. benthamiana*, we found proteins effectively inhibited aphid survival and growth. Coupled with transgenic approaches these three SerPIN-IIs are a new resource that can be further developed for the control of aphids and other insect pests.

## Acknowledgement

We thank Kerry Pedley, Jen Brady, and Greg Martin for providing the VIGS plants for aphid screening bioassays; and we thank Kim Hall and Sarah Cohen for their assistance with performing the aphid screening bioassays.

---

**Figure S1.**
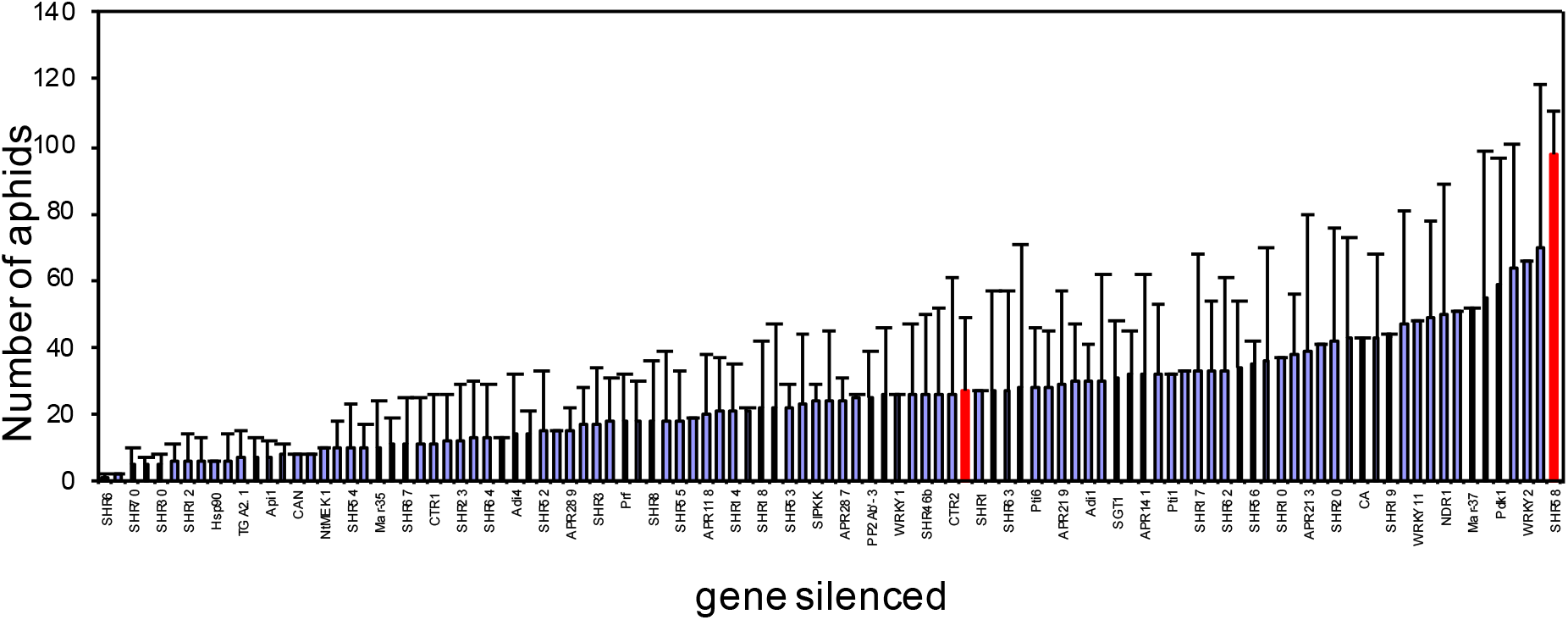
Aphid bioassays on *Nicotiana benthamiana* VIGS plants. Each bar represents the average percentage of aphid survival +/- s.e. of n = 14 for the empty vector control and n = 8 for VIGS plants. The empty vector control and SHR58 (PIN-II) are highlighted in red.

**Figure S2.**
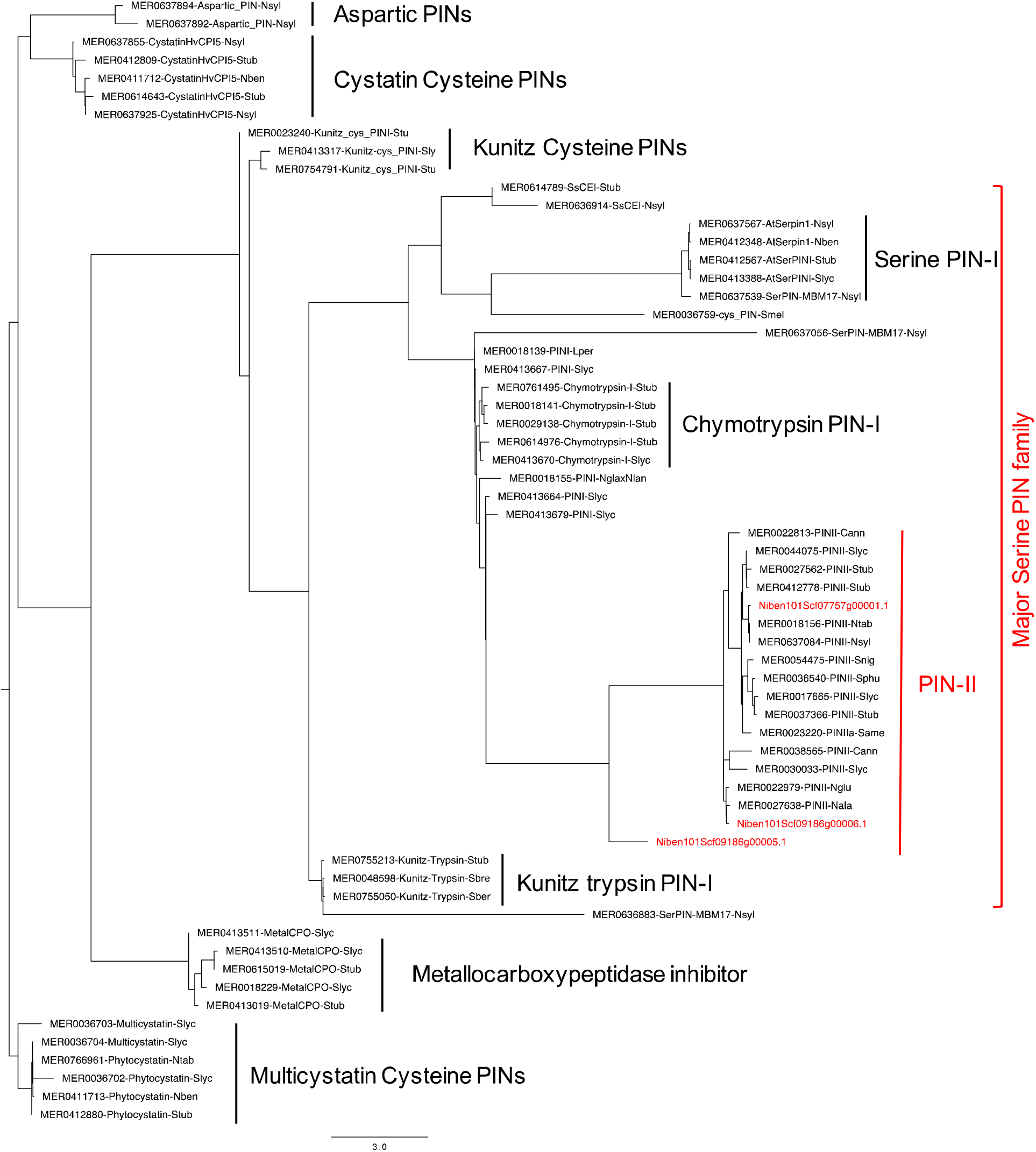
Phylogeny of proteinase inhibitors from the Solanaceae family. A maximum likelihood tree was built using RAxML with 1000 bootstraps. PINs from all four major families were included, *i.e.,* cysteine PINs, metalloid PINs, aspartic PINs, and serine PINs. The three *N. benthamiana* PINs identified in this study, which were clustered in a mono-phylogenetic PIN-II group within the major serine PIN family, are highlighted in red. The naming of each protein (*e.g.,* MER0412880-Phytocystatin-Stub) includes the MER####### IDs from the MEROPS database, followed by the PIN name (Phytocystatin), and the originated plant species (Stub). The Solanaceae plant species collected included, Cann: *Capsicum annuum*; Lper: *Lycopersicon peruvianum*; Nala: *Nicotiana alata*; Nben: *N. benthamiana*; Ngla: *N. glauca*; Nglu: *N. glutinosa*; Nlan: *N. langsdorffii*; Ntab: *N. tabacum*; Nsyl: *N. sylvestris*; Same: *S. americanum*; Sber: *S. berthaultii*; Sbre: *S. brevidens*; Slyc: *S. lycopersicum*; Snig: *S. nigrum*; Sphu: *S. phureja*; Stub: *Solanum tuberosum*.

**Figure S3.**
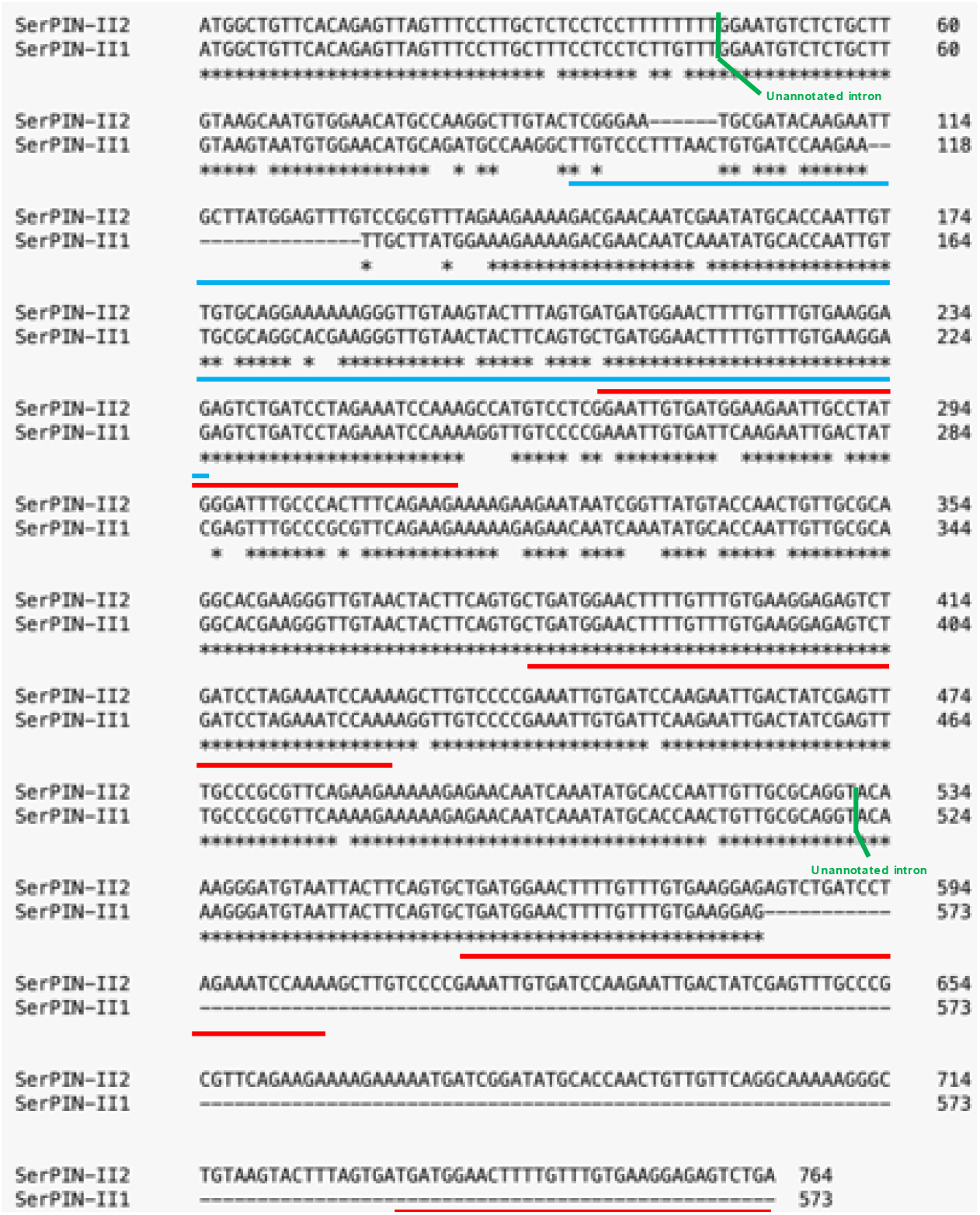
*SerPIN-II1 and SerPIN-II2* sequence analyses. The blue underlines indicate the dsRNA for the VIGS experiment (Figure 3A and S1), which is identical in *N. benthamiana SerPIN-II2* and tomato. The dsRNA likely has enough identity to also reduce expression of *SerPIN-II1*. Red underlining indicates the exact repeated regions within the coding sequences. Green bars indicate unannotated introns with unknown lengths (shown as Ns in the genome assembly).

**Figure S4.**
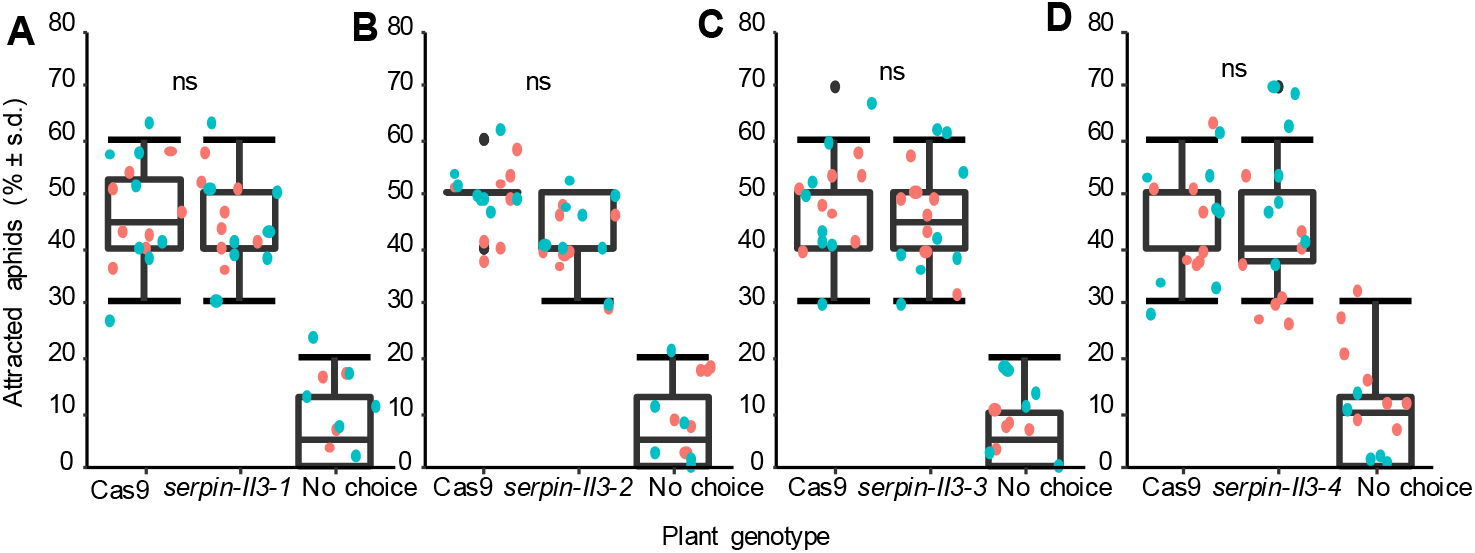
Aphid choice assays with *serpin-II3* mutants and *Cas*9-expressing control *N. benthamiana*. Aphids were exposed to detached leaves in Petri dishes for 24 h. Graphs show Cas9-expressing controls versus: (A) *serpinII3-1*; (B) *serpinII3-2*; (C) *serpinII3-3*; (D) *serpinII3-4*. Significance between genotypes was tested using Chi-squared tests. ns = not significant (P > 0.05).

